# aNy-way Independent Component Analysis

**DOI:** 10.1101/2020.02.03.930156

**Authors:** Kuaikuai Duan, Rogers F. Silva, Vince D. Calhoun, Jingyu Liu

**Author notes:** These authors have contributed equally to this work (Kuaikuai Duan, Rogers F. Silva).

## Abstract

Multimodal data fusion is a topic of great interest. Several fusion methods have been proposed to investigate coherent patterns and corresponding linkages across modalities, such as joint independent component analysis (jICA), multiset canonical correlation analysis (mCCA), mCCA+jICA, disjoint subspace using ICA (DS-ICA) and parallel ICA. JICA exploits source independence but assumes shared loading parameters. MCCA maximizes correlation linkage across modalities directly but is limited to orthogonal features. While there is no theoretical limit to the number of modalities analyzed together by jICA, mCCA, or the two-step approach mCCA+jICA, these approaches can only extract common features and require the *s*a*me* number of sources/components for all modalities. On the other hand, DS-ICA and parallel ICA can identify both common and distinct features but are limited to two modalities. DS-ICA assumes shared loading parameters among common features, which works well when links are strong. Parallel ICA simultaneously maximizes correlation between modalities and independence of sources, while allowing different number of sources for each modality. However, only a very limited number of modalities and linkage pairs can be optimized. To overcome these limitations, we propose aNy-way ICA, a new model to simultaneously maximize the independence of sources and correlations across modalities. aNy-way ICA combines infomax ICA and Gaussian independent vector analysis (IVA-G) via a shared weight matrix model without orthogonality constraints. Simulation results demonstrate that aNy-way ICA not only accurately recovers sources and loadings, but also the true covariance/linkage patterns, whether different modalities have the same or different number of sources. Moreover, aNy-way ICA outperforms mCCA and mCCA+jICA in terms of source and loading recovery accuracy, especially under noisy conditions.

**Clinical Relevance:** This establishes a model for N-way data fusion of any number of modalities and linkage pairs, allowing different number of non-orthogonal sources for different modalities.

## I. Introduction

With the fast advancement of technology, growing medical data from different imaging modalities and omics (e.g. genomics, transcriptomics, epigenomics, microbiomics) can be collected from the *same* participants for a variety of diseases. Data from different modalities/omics provide unique and complementary features for understanding diseases. Thus, integrative/fusion methods are essential to identify unique and coherent biomarkers across modalities.

Several fusion approaches have been proposed to explore the latent shared structures in multimodal datasets in order to discover and establish links among modalities. To that end, joint independent component analysis (jICA) [1] has focused on identifying shared loadings across modalities. Meanwhile, mCCA+jICA [2] proposed a multiset canonical correlation analysis (mCCA) preprocessing in order to improve the correspondence over modalities and more closely satisfy the assumption of shared identical loadings prior to jICA, enabling N-way multimodal fusion. With similar motivation, disjoint subspace using ICA (DS-ICA) [3] proposed to apply jICA on the common subspace extracted with principal component analysis-CCA (PCA-CCA) [4], and separate ICAs on the remaining (distinct) subspace. Parallel ICA [5], on the other hand, proposed to interweave separate (parallel) ICAs, one per modality, with simultaneous maximization of correlation between specific multimodal loading pairs. This is attractive because both independence and linkage are optimized together rather than in two steps. However, Parallel ICA (and 3-way Parallel ICA) [5, 6] is limited in the number of modalities and linked components it can robustly detect since it optimizes individual correlation pairs rather than the entire underlying linkage structure.

Here, we propose a new model called aNy-way ICA, which overcomes these limitations. aNy-way ICA optimizes the entire loadings correlation structure of linked components via Gaussian independent vector analysis (IVA-G) [7] and simultaneously optimizes independence via separate (parallel) ICAs. Therefore, aNy-way ICA is capable of detecting multiple linked sources over any number of modalities without requiring orthogonality constraints on sources and permitting different number of sources for different modalities. The remainder of the paper is organized as follows. Section II introduces the aNy-way ICA model. Section III describes simulation scenarios. Results and conclusions are shown in sections IV and V, respectively.

## II. METHOD

aNy-way ICA aims to simultaneously maximize the independence of sources ***s*** while minimizing the mutual information between subspace component vectors (SCV) **a**_*k*_. As shown in Fig. 1, without loss of generality, we consider three modalities: structural MRI (sMRI), functional MRI (fMRI) and electroencephalography (EEG), each with 5, 4, and 6 sources, respectively. While the total number of sources *C*_*m*_ is different per modality, at most *K* = 4 SCVs (maximum number of one-to-one correspondences across all modalities) can be extracted. For each modality, ICA decomposes the data **X**_*m*_ (dimension: *N* × *L*_*m*_; *m* = 1,2,3 modality indices; *N* is the number of subjects; *L*_*m*_ is the feature dimensionality) into a source matrix **S**_m_ (dimension: *C*_*m*_ × *L*_*m*_) with corresponding loading matrices **A**_*m*_ (dimension: *N* × *C*_*m*_), i.e., **X**_*m*_ = **A**_*m*_**S**_*m*_.

**Figure 1.**
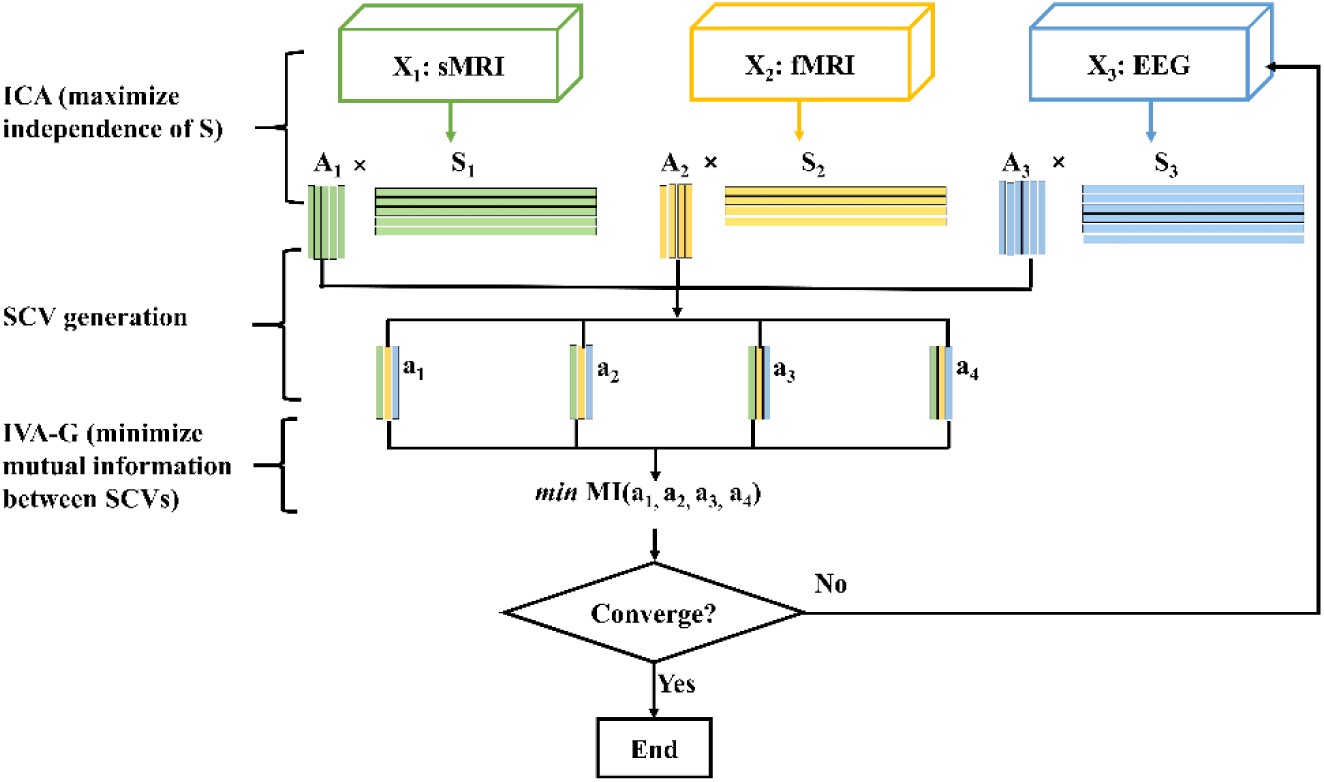
Diagram of aNy-way ICA. Sources **S**_*m*_ are estimated by maximization of independence in each modality separately. Corresponding loadings **A**_*m*_ are then organized into SCVs **a**_*k*_, followed by minimization of their mutual information with IVA-G, which amounts to maximization of the correlation structure within the SCV without orthogonality constraints [8]. The model is optimized with stochastic gradient descent until convergence.

The source matrix is estimated by the product of a weight matrix **W**_*m*_ and data as **S**_*m*_ = **W**_*m*_**UX**_*m*_, where **U** is a fixed projection into a *shared* variance-adjusted space (see below). The loading matrix is then reconstructed as the product of data and pseudo-inverse of the source matrix 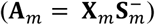, a key strategy of this work. SCVs are then formed by concatenation of linked loadings across the three modalities, where the total number of SCVs *K* is the minimum component number among the three modalities. Assuming each SCV follows a multivariate Gaussian distribution, mutual information among SCVs is minimized utilizing IVA-G since that has been shown to correspond to mCCA’s generalized variance (GENVAR) approach without orthogonality constraints [8].

The cost function o^*m*^f aNy-way ICA is thus defined as:

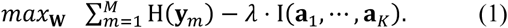

where *M* is the total number of modalities, H(**y**_*m*_) represents the entropy of **y**_*m*_ (the same as Infomax ICA cost [9]), **y**_*m*_ is a nonlinear transformation of the bias-adjusted source **u**_*m*_ 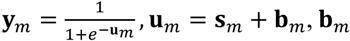 is a bias), I(**a**_1_, …, **a**_*K*_) is the mutual information among SCVs **a**_*k*_, *λ* is a regularizer to balance between independence (ICA) and correlation (IVA-G) maximization. The entropy of **y**_*m*_ [9] and mutual information among SCVs **a**_*k*_ [7] are defined in (2) and (3), respectively:

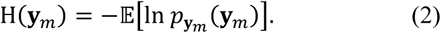

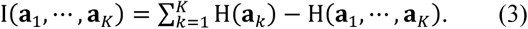

where 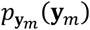 is the probability density function of the vector **y**_*m*_. The *k*-th SCV **a**_*k*_ can be formed as 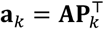, where 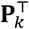 is akin to a permutation matrix that selects the loadings from the *k*-th component of each modality, and **A** = [**A**_1_, …, **A**_*M*_] is a concatenation of loadings from all modalities.

H(**a**_*k*_) represents the marginal entropy of the *k*-th SCV and H(**a**_1_, …, **a**_*K*_) 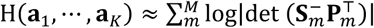 is the joint entropy of all SCVs, which simplifies to a determinant function on the sources after noting the entropy of the data is constant. The proposed aNy-way ICA is then solved by stochastic gradient descent (SGD), utilizing the relative gradients [10] of (2) and (3) with respect to **W**, which are given, respectively, as:

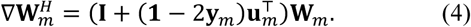

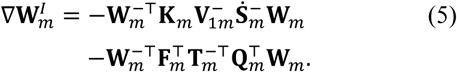

where **I** is an identity matrix, **1** is a column vector of ones, ⊤ is the matrix transpose, 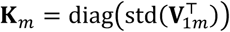 is a diagonal matrix, and 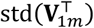 is a vector with the standard deviations from each row of 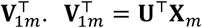 is based on the eigenvalue decomposition (EVD) of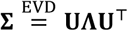, with 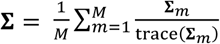 being the average of normalized sample covariance matrices from each modality 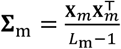 (**X**_*m*_ has zero mean rows and columns). 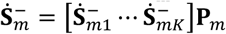**P***m* is an intermediate gradient for the pseudo-inverse of the source matrix of the *m*-th modality, and 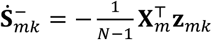 with 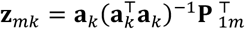. Note that 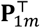 selects the *m*-th column from the matrix to its left, and **P**_*m*_ expands the concatenation of 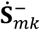 vectors from *L*_*m*_ × *K* to *L*_*m*_ × *C*_*m*_ (effectively inserting zero columns as needed). Finally, approximating the determinants in the SCV joint entropy term as the product of singular values of 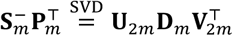 (via singular value decomposition (SVD)) and collecting constant terms into 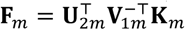 and 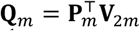, yields the final form in (5), where 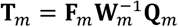.

Since the source/loading ordering from separate ICAs per modality would be arbitrary (not linked), it is important *not* to over-emphasize independence early during the optimization. Rather, we propose to project the multimodal data into a *shared* variance-adjusted space, i.e., along the shared basis **U** defined above, which provides a good (linked) starting point for the optimization. Also, we observed improved loading (SCV) linkage and source alignment by initially optimizing only IVA-G for a few steps/epochs (here, 5) before optimizing (1). Moreover, the regularizer parameter *λ* is initialized with 1 and adaptively adjusted by annealing it according to the entropy trend during the optimization: whenever entropy decreases with a slope larger than a threshold (here, 10^−4^), this triggers *λ* to be reduced by a factor (here, 0.8).

## III. SIMULATION

Simulated sMRI, fMRI and EEG data were employed to validate the proposed aNy-way ICA, where sMRI and fMRI sources were simulated with the SimTB toolbox [11]. For EEG sources, we used a subset of simulated independent Laplacian timeseries in [12], which had an autocorrelation around 0.85 between timepoint *t* and *t* − 1, and 0.2 between *t* and *t* − 10. Each SCV was sampled from a zero mean *m*-dimensional multivariate Gaussian distribution with unique covariance structures (correlations’ range: [0.2,0.8]). SCVs were then organized into loading matrices **A**_*m*_. If applicable, extra loadings for non-linked sources were sampled from a standard Normal distribution and padded to **A**_*m*_. Finally, the data was generated as the product of the loading and source matrices as **X**_*m*_ = **A**_*m*_**S**_*m*_. The following scenarios were then examined:

### Scenario 1

All 3 modalities had the same number of sources *C*_*m*_ = 7. The number of subjects was fixed at *N* = 500. Feature dimensions (*L*_*m*_) for sMRI, fMRI and EEG data were 31064 (voxels), 17420 (voxels), 4444 (time points), respectively (same *L*_*m*_ used for scenario 2). Comparisons were performed against both mCCA and mCCA+jICA.

### Scenario 2

sMRI, fMRI, and EEG data had different number of sources. In part (a), we assessed the effect of varying the number of linked SCVs (*K* = [3,5,7,9,11]) while keeping *N* = 500 fixed. The number of sources *C*_*m*_ for sMRI, fMRI and EEG data were set as *K* + 1, *K*, and *K* + 2, respectively, except for *K* = 11 where *C*_1_ = *C*_2_ = 11, and *C*_3_ = 12. In part (b), the number of sources *C*_*m*_ for sMRI, fMRI and EEG data were fixed at 8, 7, 9, respectively, while the subject number *N* varied from 100 to 1000. For each subject number, 10 sets of SCV/loadings were generated and run for each modality. Finally, in part (c), we assessed the effect of additive Gaussian white noise with *C*_*m*_ fixed at 8, 7, 9 for sMRI, fMRI and EEG, respectively, and *N* fixed at 500. The signal-to-noise ratio (SNR) varied from 0db to 30dB with a step size of 5dB. For each SNR level, the noise samples **E**_*im*_ were generated 5 times, yielding noisy data 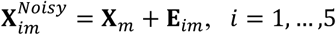, *i* = 1, …, 5 for modality *m*. Note: scenarios 1, 2(a), and 2(b) were noise-free.

The following component-wise performance measures were computed per modality for aNy-way ICA, mCCA and mCCA+jICA: correlations between recovered *sources* and ground-truth, as well as correlations between recovered *loadings* and ground-truth. To examine SCV recovery, we also calculated the MSE value between the recovered and ground-truth SCV cross-correlations (linkages), considering the diagonal blocks and off-diagonal portions separately, as well as together (whole matrix). For mCCA and mCCA+jICA, we estimated both 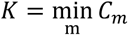 and 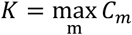 for all modalities (two separate runs). For scenario 2(a), correlations were averaged across matched components per modality, while for scenarios 2(b) and 2(c), correlations were averaged across runs and matched components. MSE values were averaged across runs.

## IV. RESULT

Fig. 2 illustrates the seven sources for (a) sMRI, (b) fMRI, and (c) EEG data in scenario 1. Fig. 3 (a)-(d) shows their recovery accuracies: under noise-free conditions, aNy-way ICA recovered all designed SCV structures very well (the MSE value was 2.8E-4). Both mCCA and mCCA+jICA failed to recover, as indicated by the large values on off-diagonal positions, likely due to overfitting of correlations and inability to estimate non-orthogonal sources in mCCA, plus potential residual source misalignments from mCCA leaking into jICA for mCCA+jICA. Fig. 3 (e)-(f) demonstrates aNy-way ICA accurately recovered both sources and loading coefficients for all three modalities, contrary to mCCA and mCCA+jICA.

**Figure 2.**
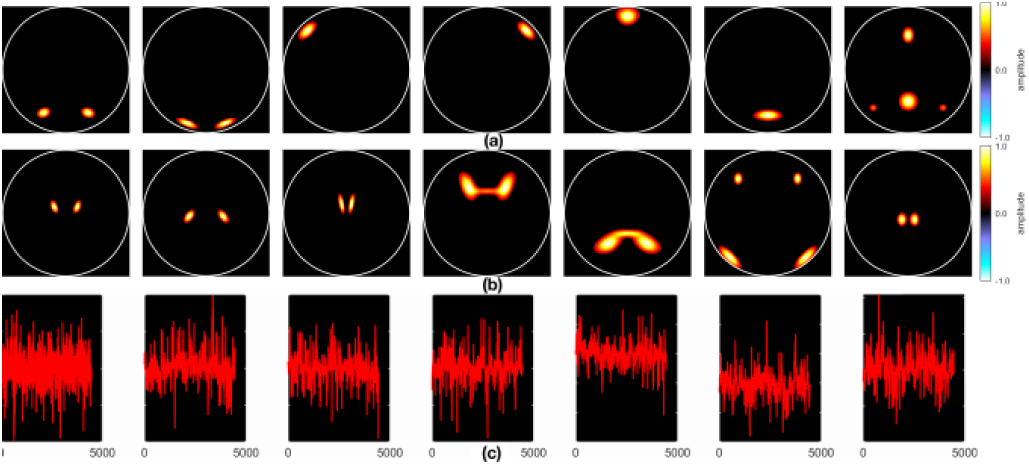
*Scenario 1* sources for (a) sMRI, (b) fMRI, and (c) EEG data.

**Figure 3.**
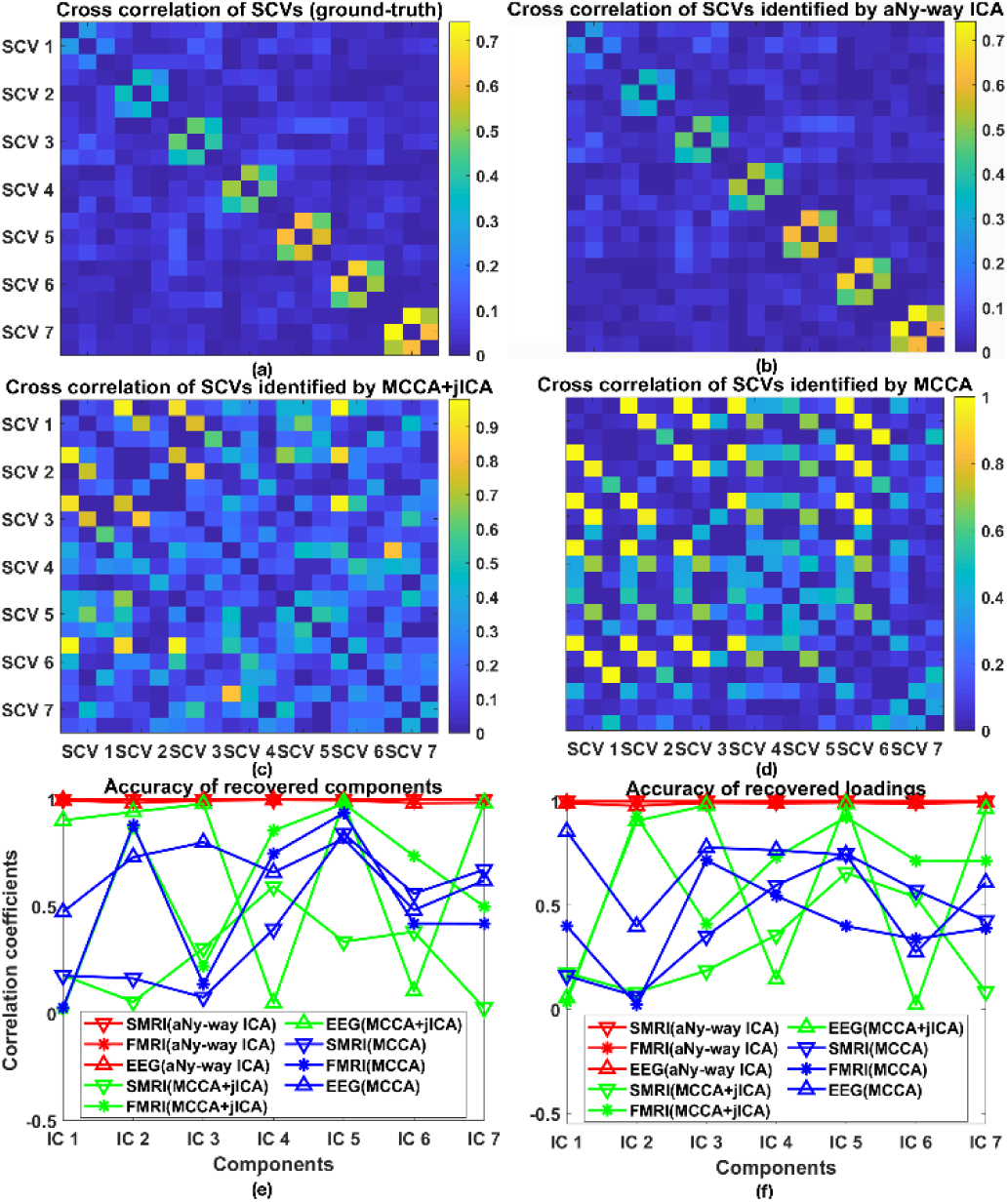
*Scenario* 1. Cross-correlation of SCVs from (a) groundtruth, (b) aNy-way ICA, (c) mCCA+jICA, and (d) mCCA. Correlation coefficients between (e) recovered sources and ground-truth, and between (f) recovered loadings and ground-truth for each independent component (IC): sMRI (downward triangle), fMRI (star), and EEG data (upward triangle) from aNy-way ICA (red), mCCA+jICA(green) and mCCA(blue). Note that shapes and colors are consistent throughout the paper.

Performance measures for scenario 2(a) are shown in Fig. 4. aNy-way ICA accurately recovered components and loadings for sMRI and fMRI data, with slightly reduced recovery accuracy for EEG data (especially for 9 and 11 SCVs). The inter-symbol interference [13] for the recovered loading matrix and minimum distance indices [14] for the recovered source matrix corroborate these observations and, thus, are omitted here. Lower accuracies for EEG data are likely due to its relatively small feature dimensionality (*L*_3_ = 4444) and weaker designed correlation linkages with other modalities within each SCV. Overall, aNy-way ICA outperformed mCCA+jICA and mCCA in terms of source and loading recovery. mCCA+jICA and mCCA decompositions with the minimum component number of the three modalities had higher accuracies compared to those obtained with the maximum component number. As the SCV number increased (e.g., 9 or 11 SCVs), source and loading accuracies from both mCCA and mCCA+jICA dropped. Table I lists MSE values for aNy-way ICA’s recovered covariance structure. aNy-way ICA fully recovered the designed covariance structure when SCV number was 3, 5, and 7, and largely recovered it when SCV number was 9 and 11 (actually, sMRI and fMRI were aligned well but not sMRI-EEG and fMRI-EEG).

**Figure 4.**
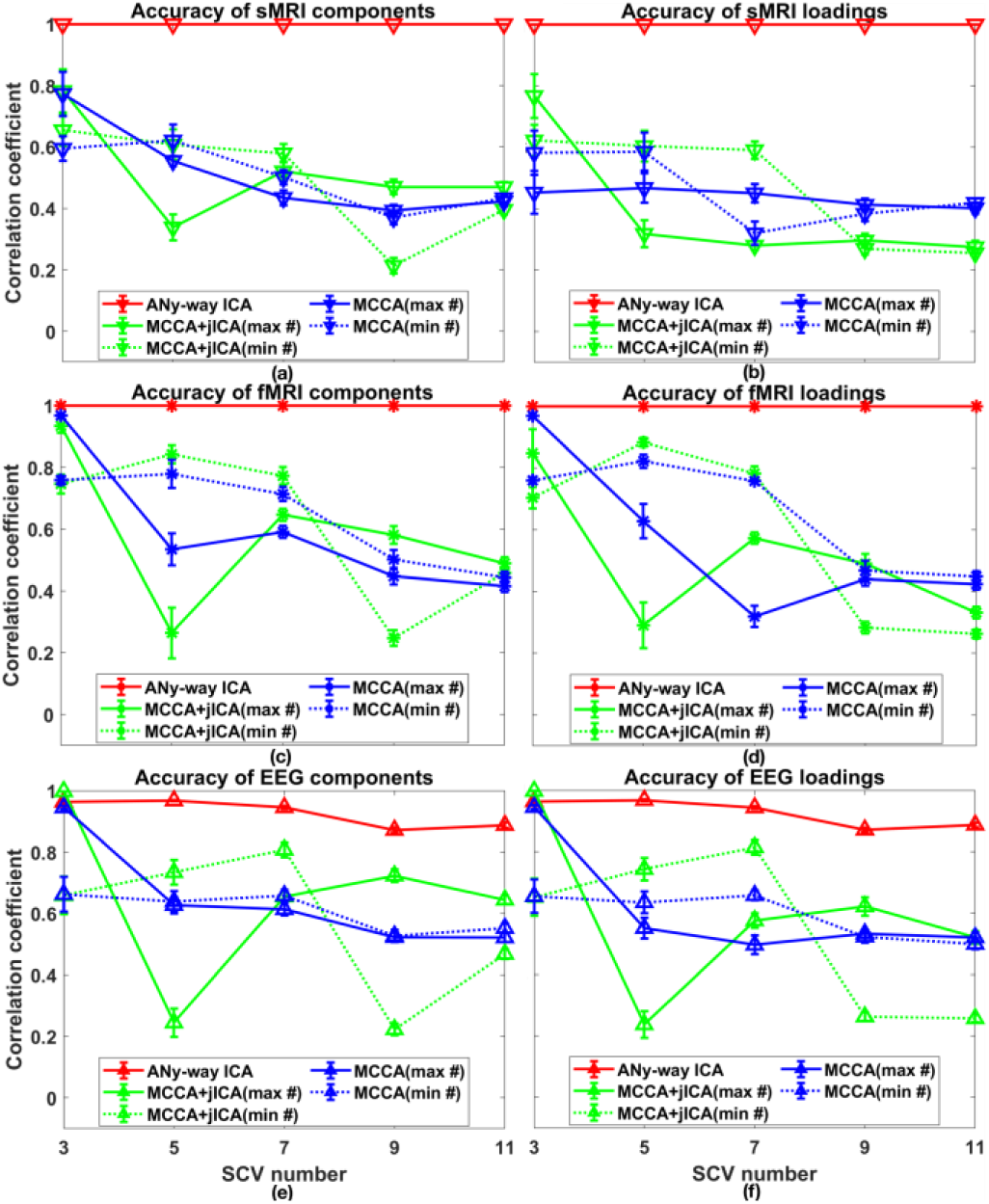
*Scenario 2(a)*, where the number of the SCVs varied from 3 to 11. Recovered *source* accuracies for (a) sMRI, (c) fMRI, and (e) EEG data, and *loading* accuracies for (b) sMRI, (d) fMRI, and (f) EEG data using aNy-way ICA (red), mCCA+jICA with maximum (solid green) or minimum component number (dashed green), mCCA with maximum (solid blue) or minimum component number (dashed blue). Values are averaged across matched components per modality.

**TABLE I.** MSE Values Between the Recovered and Ground-Truth SCV Cross-Correlations by aNy-Way ICA while Varying the Number of SCVs

| MSE | 2(a) Varying Number of SCVs ($K$) | | | | |
| --- | --- | --- | --- | --- | --- |
|  | 3 | 5 | 7 | 9 | 11 |
| All* | $2 \times 10^{-4}$ | $5 \times 10^{-4}$ | $6 \times 10^{-4}$ | $2 \times 10^{-2}$ | $6 \times 10^{-3}$ |
| Diag* | $3 \times 10^{-5}$ | $2 \times 10^{-5}$ | $2 \times 10^{-4}$ | $8 \times 10^{-2}$ | $3 \times 10^{-2}$ |
| Off* | $3 \times 10^{-4}$ | $7 \times 10^{-4}$ | $6 \times 10^{-4}$ | $9 \times 10^{-3}$ | $3 \times 10^{-3}$ |

\*Note: 'All' denotes MSE of whole matrix, 'Diag' denotes MSE of diagonal blocks only, 'Off' denotes MSE of off-diagonal blocks only.

Fig. 5 demonstrates performance measures for scenario 2(b). As number of subjects increased, the recovery accuracies of sources and loadings for fMRI and EEG, as well as the covariance structure of SCVs, increased, although there were some variations when the subject number was less than 400. Variation of accuracies of EEG data from aNy-way ICA were probably due to weak covariance structures generated from few samples. MSE values for the whole matrix, diagonal blocks, and off-diagonal blocks of the recovered SCV cross-correlation are listed in Table II 2(b). As the number of subjects increased, the MSE values decreased (except for *N* = 400 and *N* = 1000).

**Figure 5.**
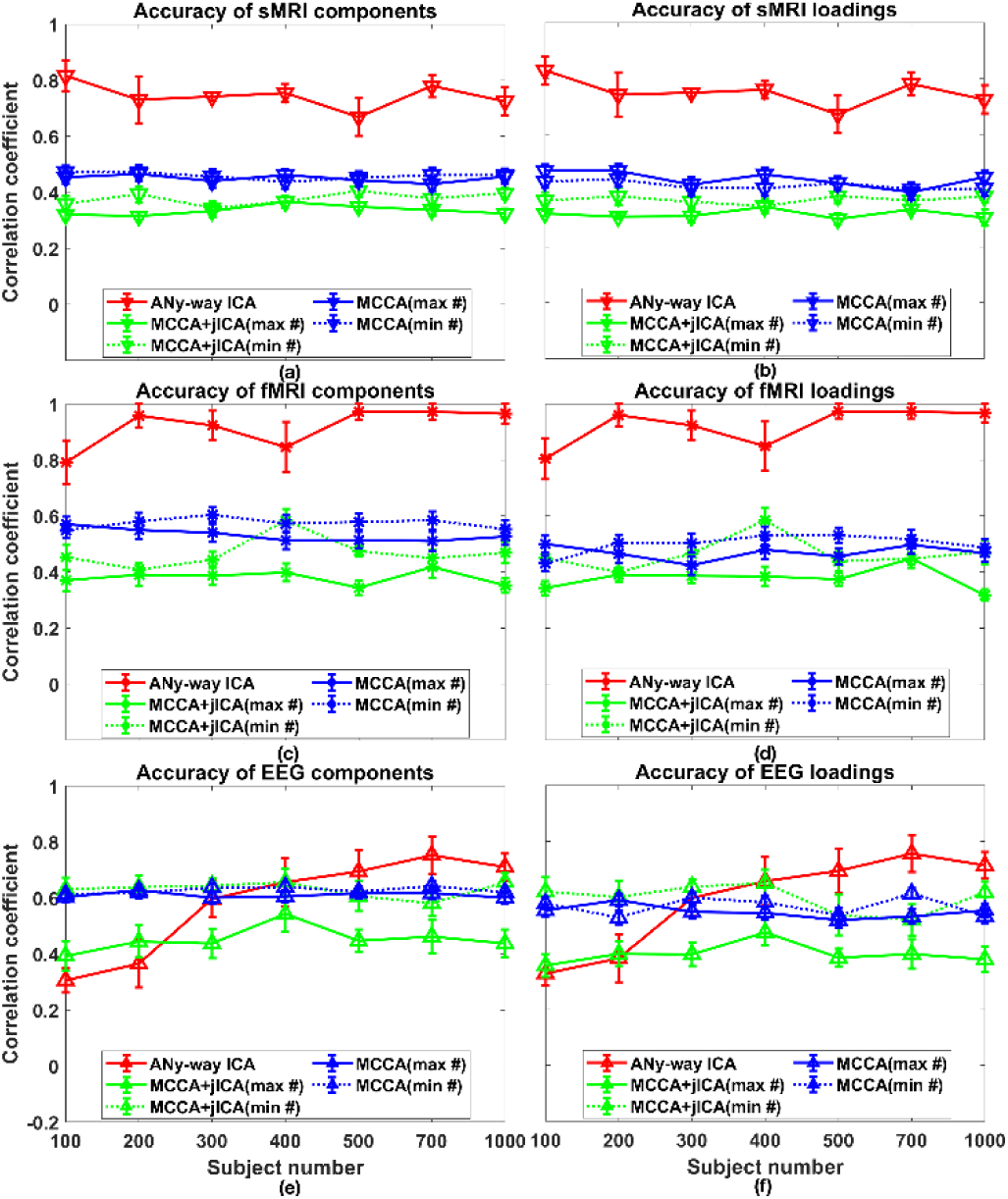
*Scenario 2(b)*, where the number of subjects *N* varied from 100 to 1000. Recovered *source* accuracies for (a) sMRI, (c) fMRI, and (e) EEG data, and *loading* accuracies for (b) sMRI, (d) fMRI, and (f) EEG data. Values are averaged across runs and matched components per modality.

**TABLE II.** MSE Values Between Recovered and Ground-Truth SCV Cross-Correlations by aNy-Way ICA while Varying the Number of Subjects or SNR.

| MSE |  |  |  |  |  |  |  |
| --- | --- | --- | --- | --- | --- | --- | --- |
| 2(b) Varying Number of Subjects (N) |  |  |  | 2(c) Varying SNR |  |  |  |
| N | All* | Diag* | Off* | SNR | All* | Diag* | Off* |
| 100 | $2 \times 10^{-2}$ | $5 \times 10^{-2}$ | $1 \times 10^{-2}$ | 0 | $4 \times 10^{-2}$ | $2 \times 10^{-1}$ | $1 \times 10^{-2}$ |
| 200 | $1 \times 10^{-2}$ | $4 \times 10^{-2}$ | $8 \times 10^{-3}$ | 5 | $3 \times 10^{-2}$ | $2 \times 10^{-1}$ | $4 \times 10^{-3}$ |
| 300 | $8 \times 10^{-3}$ | $2 \times 10^{-2}$ | $5 \times 10^{-3}$ | 10 | $3 \times 10^{-2}$ | $1 \times 10^{-1}$ | $1 \times 10^{-2}$ |
| 400 | $1 \times 10^{-2}$ | $3 \times 10^{-2}$ | $6 \times 10^{-3}$ | 15 | $1 \times 10^{-2}$ | $7 \times 10^{-2}$ | $2 \times 10^{-3}$ |
| 500 | $8 \times 10^{-3}$ | $3 \times 10^{-2}$ | $5 \times 10^{-3}$ | 20 | $8 \times 10^{-3}$ | $3 \times 10^{-2}$ | $4 \times 10^{-3}$ |
| 700 | $5 \times 10^{-3}$ | $1 \times 10^{-2}$ | $4 \times 10^{-3}$ | 25 | $2 \times 10^{-3}$ | $5 \times 10^{-3}$ | $1 \times 10^{-3}$ |
| 1000 | $6 \times 10^{-3}$ | $2 \times 10^{-2}$ | $4 \times 10^{-3}$ | 30 | $9 \times 10^{-4}$ | $4 \times 10^{-4}$ | $1 \times 10^{-3}$ |

\*Note: 'All' denotes MSE of whole matrix, 'Diag' denotes MSE of diagonal blocks only, 'Off' denotes MSE of off-diagonal blocks only.

Performance measures for scenario 2(c) are plotted in Fig. 6. Increasing the strength of noise superimposed on data, recovery accuracies of sources and loadings dropped. aNy-way ICA outperformed mCCA and mCCA+jICA when SNR ≥ 5dB. When SNR = 0db, aNy-way ICA, mCCA, as well as mCCA+jICA failed to recover the sources and loadings. Table II 2(c) summarizes MSE values of the whole matrix, diagonal blocks and off-diagonal blocks of the recovered SCV cross-correlation, reflecting larger MSE values for smaller SNR (i.e. heavier noise).

**Figure 6.**
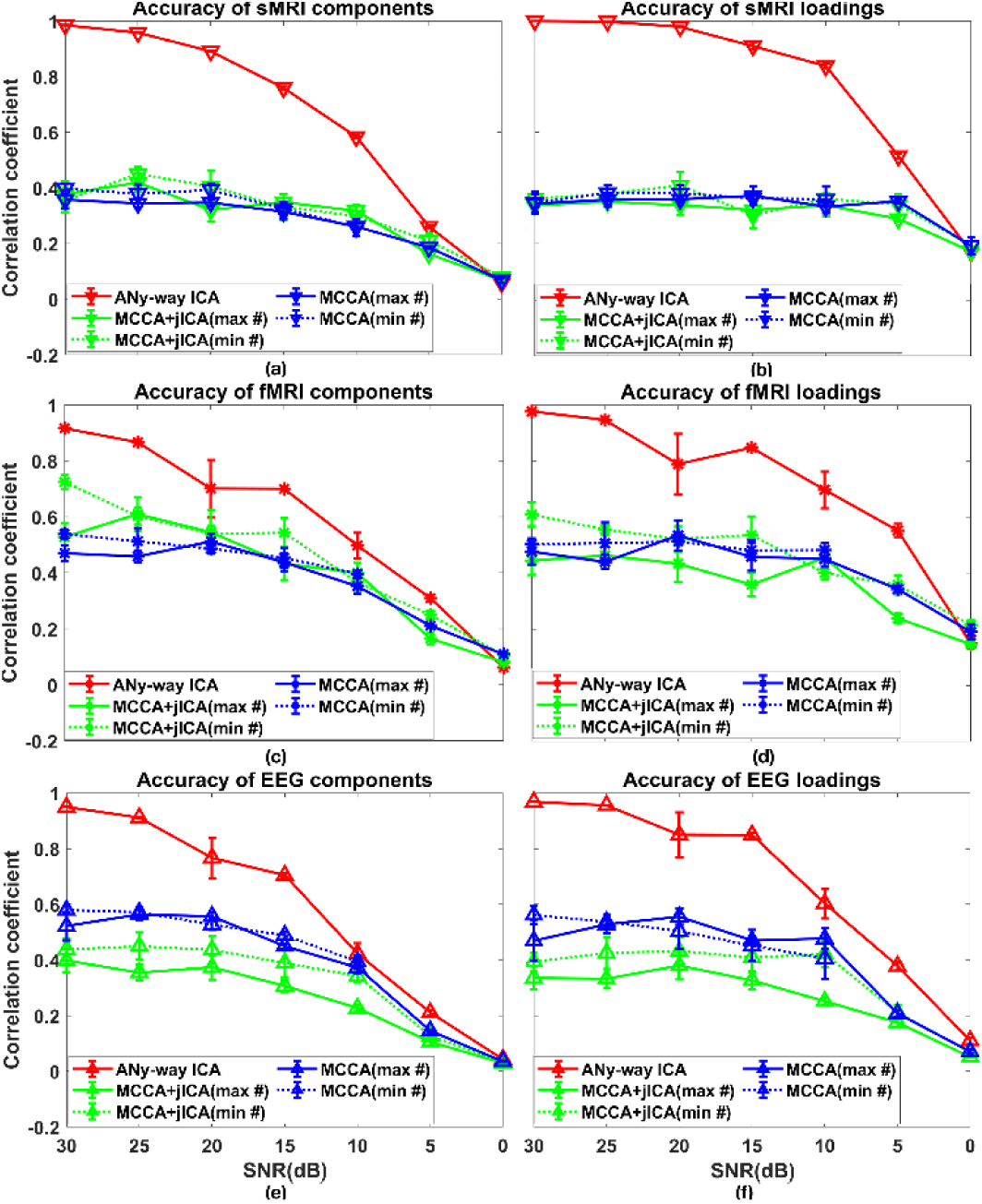
*Scenario 2(c)*, where SNR varied from 30db to 0db. Recovered *source* accuracies for (a) sMRI, (c) fMRI, and (e) EEG data, and *loading* accuracies for (b) sMRI, (d) fMRI, and (f) EEG data. Values are averaged across runs and matched components per modality.

## V. Conclusion

The proposed aNy-way ICA simultaneously optimizes independence of sources and correlations across modalities without orthogonality constraints. It can be flexibly applied to N-way multimodal fusion with the same or different number of sources per modality. Simulation results support that aNy-way ICA can recover sources and loadings, as well as the true covariance patterns, with improved accuracy compared to mCCA and mCCA+jICA, especially under noisy conditions.

## References

[1] M. Moosmann, T. Eichele, H. Nordby, K. Hugdahl, and V. D. Calhoun, “Joint independent component analysis for simultaneous EEG-fMRI: principle and simulation,” Int J Psychophysiol, vol. 67, no. 3, pp. 212–21, Mar 2008, doi: 10.1016/j.ijpsycho.2007.05.016.

[2] J. Sui et al., “Discriminating schizophrenia and bipolar disorder by fusing fMRI and DTI in a multimodal CCA+ joint ICA model,” Neuroimage, vol. 57, no. 3, pp. 839-55, Aug 1 2011, doi: 10.1016/j.neuroimage.2011.05.055.

[3] T. Adali, M. Akhonda, and V. D. Calhoun, “ICA and IVA for Data Fusion: An Overview and a New Approach Based on Disjoint Subspaces,” IEEE Sens Lett, vol. 3, no. 1, Jan 2019, doi: 10.1109/LSENS.2018.2884775.

[4] Y. Levin-Schwartz, Y. Song, P. J. Schreier, V. D. Calhoun, and T. Adali, “Sample-poor estimation of order and common signal subspace with application to fusion of medical imaging data,” Neuroimage, vol. 134, pp. 486-493, Jul 1 2016, doi: 10.1016/j.neuroimage.2016.03.058.

[5] J. Y. Liu, G. Pearlson, A. Windemuth, G. Ruano, N. I. Perrone-Bizzozero, and V. Calhoun, “Combining fMRI and SNP Data to Investigate Connections Between Brain Function and Genetics Using Parallel ICA,” (in English), Hum Brain Mapp, vol. 30, no. 1, pp. 241–255, Jan 2009, doi: 10.1002/hbm.20508.

[6] V. M. Vergara, A. Ulloa, V. D. Calhoun, D. Boutte, J. Y. Chen, and J. Y. Liu, “A three-way parallel ICA approach to analyze links among genetics, brain structure and brain function,” (in English), Neuroimage, vol. 98, pp. 386–394, Sep 2014, doi: 10.1016/j.neuroimage.2014.04.060.

[7] M. Anderson, T. Adali, and X. L. Li, “Joint Blind Source Separation With Multivariate Gaussian Model: Algorithms and Performance Analysis,” (in English), Ieee T Signal Proces, vol. 60, no. 4, pp. 1672–1683, Apr 2012, doi: 10.1109/Tsp.2011.2181836.

[8] M. Anderson, X. L. Li, and T. Adali, “Nonorthogonal Independent Vector Analysis Using Multivariate Gaussian Model,” (in English), Latent Variable Analysis and Signal Separation, vol. 6365, pp. 354–361, 2010, doi: 10.1007/978-3-642-15995-4_44.

[9] A. J. Bell and T. J. Sejnowski, “An Information Maximization Approach to Blind Separation and Blind Deconvolution,” (in English), Neural Comput, vol. 7, no. 6, pp. 1129–1159, Nov 1995, doi: 10.1162/neco.1995.7.6.1129.

[10] C. J. Pierre Comon, Handbook of Blind Source Separation: Independent Component Analysis and Applications, 1st ed. The Boulevard, Langford Lane, Kidlington, Oxford, OX5 1GB, UK: Academic Press., 2010.

[11] E. B. Erhardt, E. A. Allen, Y. H. Wei, T. Eichele, and V. D. Calhoun, “SimTB, a simulation toolbox for fMRI data under a model of spatiotemporal separability,” (in English), Neuroimage, vol. 59, no. 4, pp. 4160–4167, Feb 15 2012, doi: 10.1016/j.neuroimage.2011.11.088.

[12] R. F. Silva, S. M. Plis, T. Adali, M. S. Pattichis, and V. D. Calhoun, “Multidataset Independent Subspace Analysis with Application to Multimodal Fusion,” arXiv preprint, 2019. [Online]. Available: https://arxiv.org/abs/1911.04048.

[13] S. Amari, A. Cichocki, and H. H. Yang, “A new learning algorithm for blind signal separation,” (in English), Adv Neur In, vol. 8, pp. 757–763, 1996. [Online]. Available: <Go to ISI>://WOS:A1996BG45M00107.

[14] P. Ilmonen, K. Nordhausen, H. Oja, and E. Ollila, “A New Performance Index for ICA: Properties, Computation and Asymptotic Analysis,” (in English), Latent Variable Analysis and Signal Separation, vol. 6365, pp. 229-+, 2010, doi: 10.1007/978-3-642-15995-4_29.

